# USAT: a Bioinformatic Toolkit to Facilitate Interpretation and Comparative Visualization of Tandem Repeat Sequences

**DOI:** 10.1101/2022.04.15.488513

**Authors:** Xuewen Wang, Bruce Budowle, Jianye Ge

## Abstract

Tandem repeats (TR), which are highly variable genomic variants, are widely used in individual identification, disease diagnostics and evolutionary studies. The recent advances of sequencing technologies and bioinformatic tools facilitate calling TR haplotypes. Both length-based and sequence-based TR alleles are used in different applications. However, sequence-based TR alleles could provide the highest precision to characterize TR haplotypes. Analysis of the differences between or among TR haplotypes, especially at the single nucleotide level, is the focus of TR haplotype characterization. In this study, we developed a Universal STR Allele Toolkit (USAT) for TR haplotype analysis, which includes allele size conversion, sequence comparison of haplotypes, figure plotting and comparison for allele distribution, and interactive visualization. An example application of USAT for analysis of the CODIS core STR loci with benchmarking human individuals demonstrated the capabilities of USAT. USAT has a user-friendly graphic interface and runs in all major computing operating systems at a fast speed with parallel computing enabled. In summary, USAT is able to facilitate the interpretation, visualization, and comparisons of TRs.

## Introduction

Genomic sequence variation between and among individuals within and between species is of genetic and practical significance. Tandem repeats (TRs), a type of genomic variation, comprise a few to hundreds of tandemly repeated sequences in the genome (Willems et al. 2017; Byrska-Bishop et al. 2021). A TR can vary in the number of repeats between species and also among individuals of the same species (Fan & Chu 2007). TRs are classified into short TRs (STRs), also known as microsatellites, and variable number tandem repeats (VNTRs) or minisatellites. In particular, STRs usually contain a repeat motif, ≤ 6 base pairs (bp) in length, are widely dispersed in the genome and compose up to ∼1-3% of most eukaryotic genomes (Frazer et al. 2009; Chaisson et al. 2015; Wang & Wang 2016). TRs were known before the genomic era and were used as genetic markers (e.g., STR markers) for decades. However, it is still a challenge to complete a genome-wide analysis of TRs and understand TR significance in living organisms due to the complexity of TRs. More recently, our understanding of TRs has been increasingly enriched with the advances of better assembled genomes, high throughput DNA sequencing technologies, and bioinformatics analyses (Kistler et al. 2017; Willems *et al*. 2017; Giesselmann et al. 2019; Bakhtiari et al. 2021; Nurk et al. 2022). With the complex variabilities and high discrimination powers, TR markers have been widely used in population genetic analyses, forensic identification, molecular breeding, and selection (Frazer *et al*. 2009; Wang & Wang 2016; Saini et al. 2018; Eichler 2019; Chiu et al. 2021; Gharesouran et al. 2021). In addition, TR variations are known to associate with neural diseases, such as Alzheimer’s, obesity, and cancers via regulating proximal gene expression (Gymrek 2017; Bakhtiari *et al*. 2021). Also, STRs are the core markers of forensic DNA applications and used in almost all forensic DNA databases, such as the FBI’s Combined DNA Index System (CODIS) database (fbi.gov 2022).

In many studies and practices, the lengths of TR alleles are used, while the detailed sequence of alleles is ignored. For example, a forensic STR allele is typically recorded as the number of repeats or length-based sizes (e.g., 10.1 for an allele comprising ten repeats plus one additional base). This operationally-defined designation is due to the limitations of traditional technologies, with which the variants of TR are detected by Sanger sequencing or by measuring the lengths of DNA fragments during separation by capillary electrophoresis (CE). TR alleles with the same length are treated as the same alleles, although they may have different sequences. Higher resolution of TR alleles can be important for a wide range of applications and currently has not been fully captured.

TR alleles can be reported as sequence variants or haplotypes using next generation sequencing (NGS) technologies with higher confidence and lower cost per base pair than traditional methods (Slatko et al. 2018). Bioinformatic tools have been developed to detect TR haplotypes from sequence datasets, such as STRait Razor (Woerner et al. 2017; King et al. 2021), HipSTR (Willems *et al*. 2017) and FDSTools (Hoogenboom et al. 2017). These software programs can detect both length-based and sequence-based alleles. Each STR haplotype (i.e., sequence) contains rich information such as the number of repeats of a basic motif, and additional point mutations such as single nucleotide polymorphism (SNP) and insertions/deletions, if present. However, it is difficult to directly or visually identify the differences between TR haplotypes due to their repetitive nature, especially for complex haplotypes. In addition, some repeat expansions of disease associated TRs may be very long and contain multiple types of variants, which could further complicate comparisons.

The latest submission requirements of CODIS (fbi.gov 2022) has begun to accept the STR haplotype sequences. A conversion between sequence-based alleles and length-based alleles (i.e., the latter being the current allele designations in the CODIS system) is needed for backward compatibility purposes. Also, in many forensic mixture cases, a mixture profile typically contains multiple STR allele haplotypes from multiple contributors, and an effective comparison between these haplotypes could facilitate deconvolution of the profile.

In this study, we developed an end-user-friendly graphic bioinformatic software, Universal STR Allele Toolkit (USAT), which provides a comprehensive set of functions to analyze and visualize TR alleles, including the conversion between length-based alleles and sequence-based alleles, nucleotide comparison of TR haplotypes, an atlas of allele distributions, interactive data filtering, data formatting, and visualization in parallel computing with a graphic user interface (i.e., no command line is needed). The latest forensic recommendations for DNA forensics (Alonso et al. 2018; Phillips et al. 2018) were followed. In general, USAT facilitates the deep analysis of TR haplotypes and TR allele interpretation. The software is able to run in the major operating systems, including Windows, MacOS, and Linux.

## Results

### Workflow of USAT

USAT takes the TR sequences in a plain text file and TR loci information in a BED file as input to calculate the length of each haplotype sequence in nucleotide base pairs (bps) and the number of repeats (allele sizes) using the equation described in the method section (Figure 1). The input of TR sequences can be easily reformatted from the output of existing tools, such as the text output from STRait Razor (Woerner *et al*. 2017; King *et al*. 2021) and FDSTools (Hoogenboom *et al*. 2017), or VCF output from HipSTR (Willems *et al*. 2017). All TR data are then displayed by USAT in an interactive table for viewing, sorting, filtering, reformatting via dragging, and saving to a result file. Interactive graphic plot(s) can be generated to show an atlas of size or length distributions for selected alleles. Multiple selected sequences can be aligned with integrated MAFFT (Nakamura et al. 2018) and visualized for TR sequence comparisons, with identity marked. USAT was programmed with Java and tested in the major operating systems, such as Windows 10 (version 21H1), MacOS (version 11.6), and Ubuntu Linux (version 20.4). All functions are integrated into a user-friendly graph interface, and only mouse clicks are needed to run all analyses (Figure 1). Overall, USAT is a user-friendly software for any end-user with minimum bioinformatic skills. A command line interface of USAT calculator for converting the sequence-based alleles to length-based alleles also is provided for software developers or other pipelines as needed.

**Figure 1.**
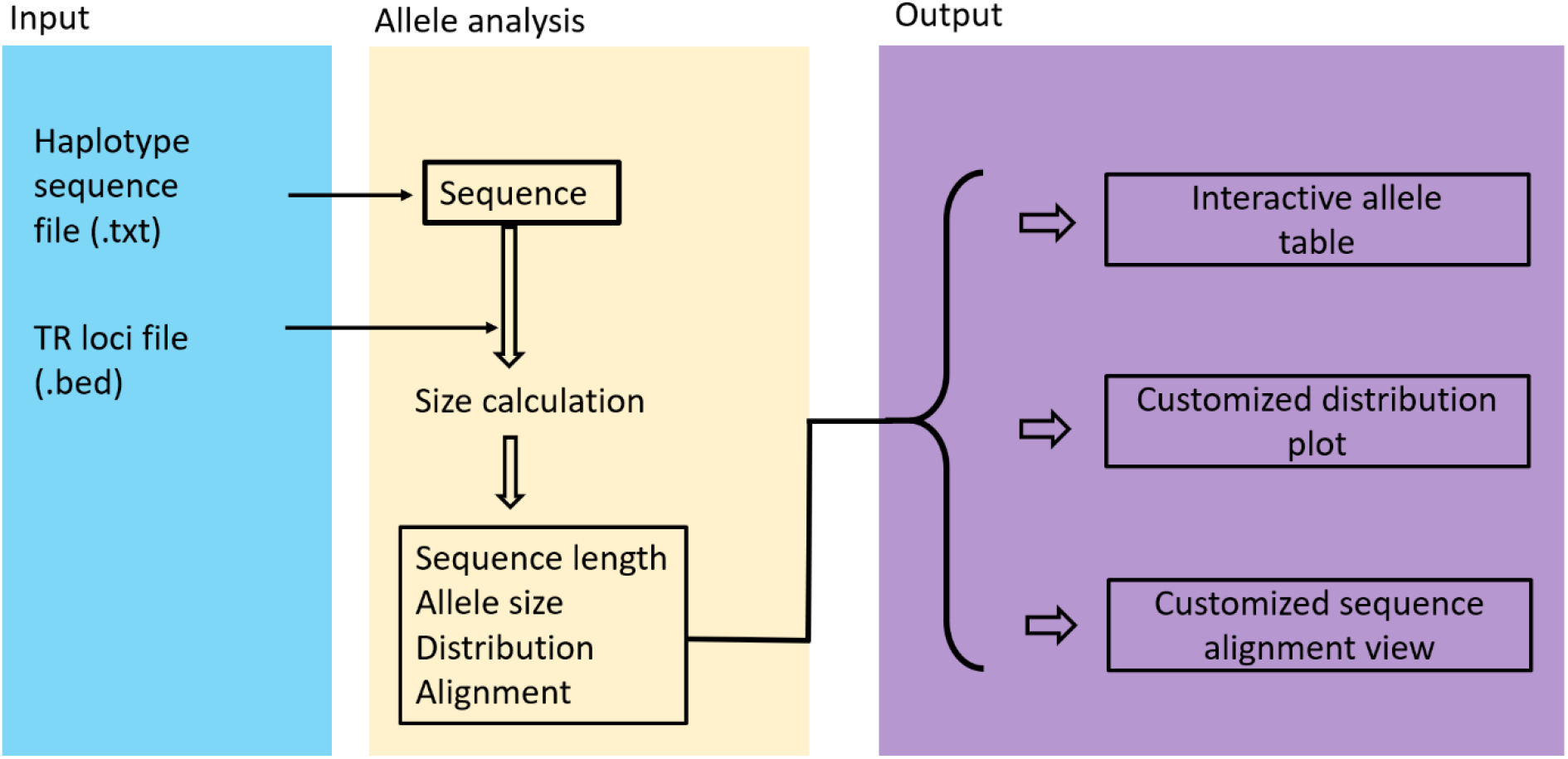
The workflow of USAT software. Three major modules of the USAT workflow are input, allele analysis, and output. The input module takes a DNA haplotype sequence file in tabular plain text format and a BED file describing the details of the tandem repeat (TR) loci. The haplotype is used to count the length in base pairs and the number of repeats (allele size) based on locus position information in the BED file. All haplotypes and calculated data are then used in displaying in an interactive table and ploting a graphic distribution requested by the user. Haplotypes of interest are aligned to identify the detailed difference between/among the haplotypes.

### Input

USAT graphic interface takes two input files in plain text format, which are the TR sequence file and a configure file for a tandem repeat locus or loci in the BED format (Figure 2). The sequence file contains a tab-separated locus or marker name, a nucleotide sequence of each TR haplotype, and sample name. One sample usually has multiple alleles (i.e., sequences), and each line is for one allele. The BED file contains the locus name, the period of motif, and the excluded length in bps for length-based forensic allele size designation. A BED file for the 20 core CODIS STR loci with coordinates of human genome assembly GRCh38 is distributed with the software.

**Figure 2.**
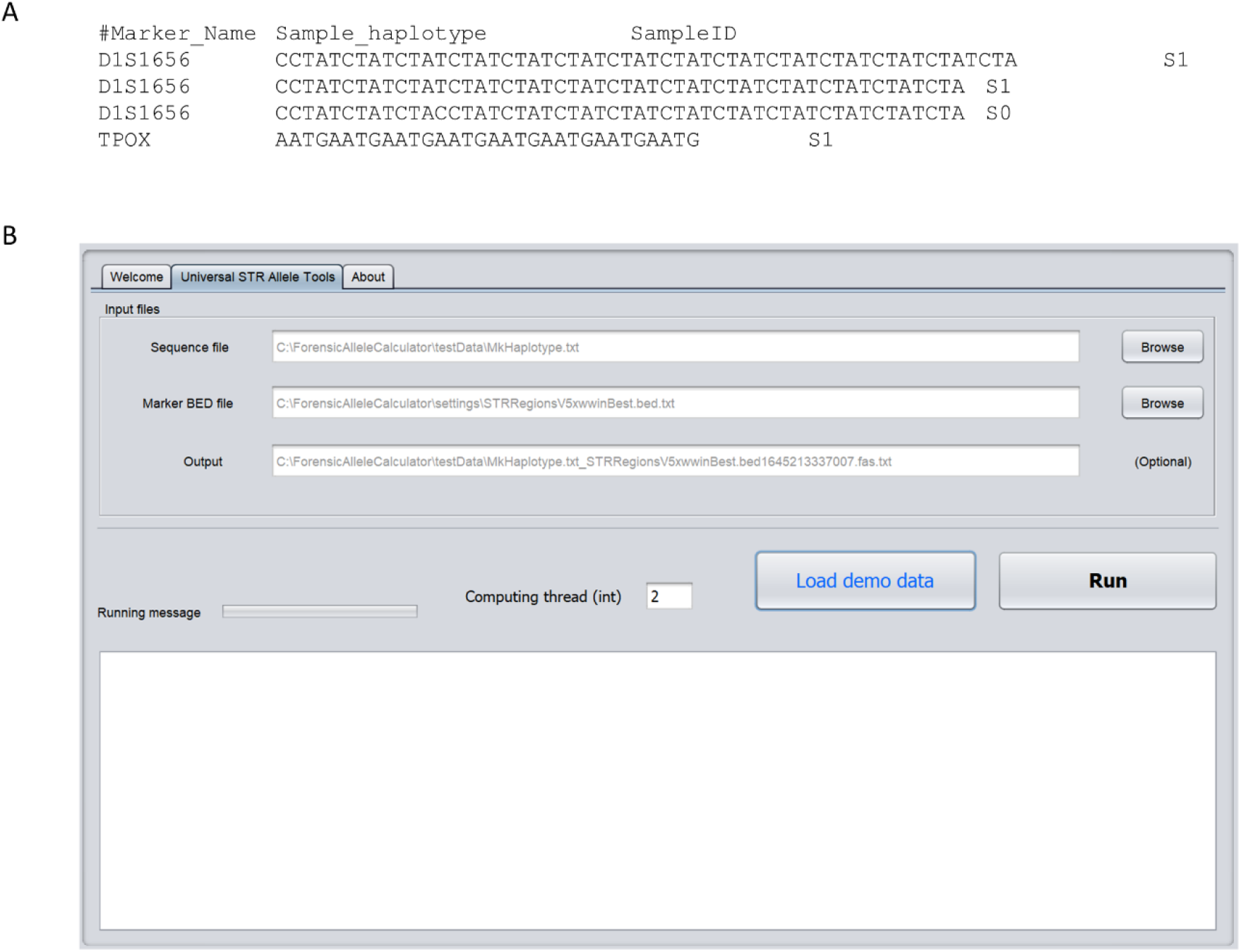
The input interface of USAT. A. An example showing the format of TR haplotype sequences in the input sequence file. B. The graphic input interface of the software USAT.

### Output

USAT reports the haplotype length, number of repeats or allele size, haplotype for each locus and sample name in an interactive table. In the table, data in each column could be sorted in ascending or descending order via just mouse clicks on the column head. The order of each column could be changed by dragging and dropping the column head. This capability could change the current format into a desired format in the output file, which can facilitate further preferred formatting for subsequent analysis (e.g., preparing for submissions to CODIS or STRidER). Figure 3 demonstrates the application of USAT for reporting alleles of the 20 CODIS core STR loci of benchmark human sample HG002 in the Genome in a Bottle (GIAB) project (https://www.nist.gov/programs-projects/genome-bottle).

**Figure 3.**
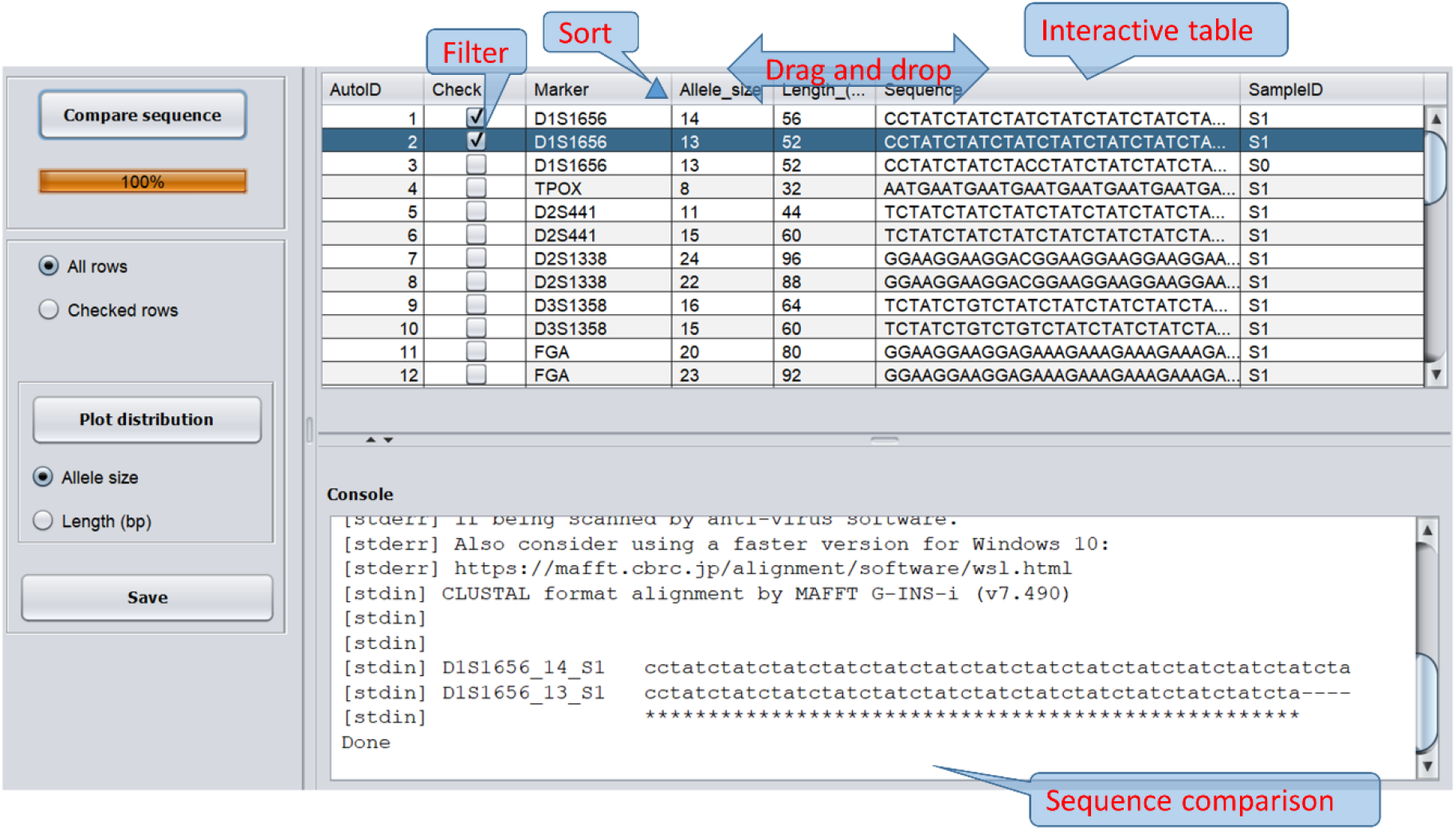
Interactive table for tandem repeat features and haplotype comparison. The top panel is an interactive table for viewing, filtering, and sorting. The bottom panel shows the sequence comparison between selected haplotypes of interest. The callouts in red letters are added as annotations for better understanding, not from the software.

To compare the haplotypic differences between multiple alleles, USAT aligns DNA sequences by calling the tool MAFFT (Nakamura *et al*. 2018) and displays the alignment by showing the difference and consistency with markers of aterisk (complete identity), colon (strong similarity), period (weak similarity), and dash for insertion/deletion, following the clustral format (www.clustal.org). Figure 3 shows the identity in alignment and the differences of two allelic sequences of marker D1S1656 of human reference HG002.

To view the atlas of allele distributions of targeted markers, interactive bar figure(s) can be plotted in USAT for any selected markers. This display enables an overviewing of the atlas of selected alleles and comparison by allele sizes or lengths. The end-user could zoom in or out of the figure and save the plot for any purpose (e.g., publication or report). For example, Figure 4 shows the atlas of a full set of 20 CODIS core loci in human sample HG002 and also the comparison of the allele size or length in bps of alleles of the selected markers. Overall, USAT provides the functions to visualize and interpret both length-based and size-based TR alleles.

**Figure 4.**
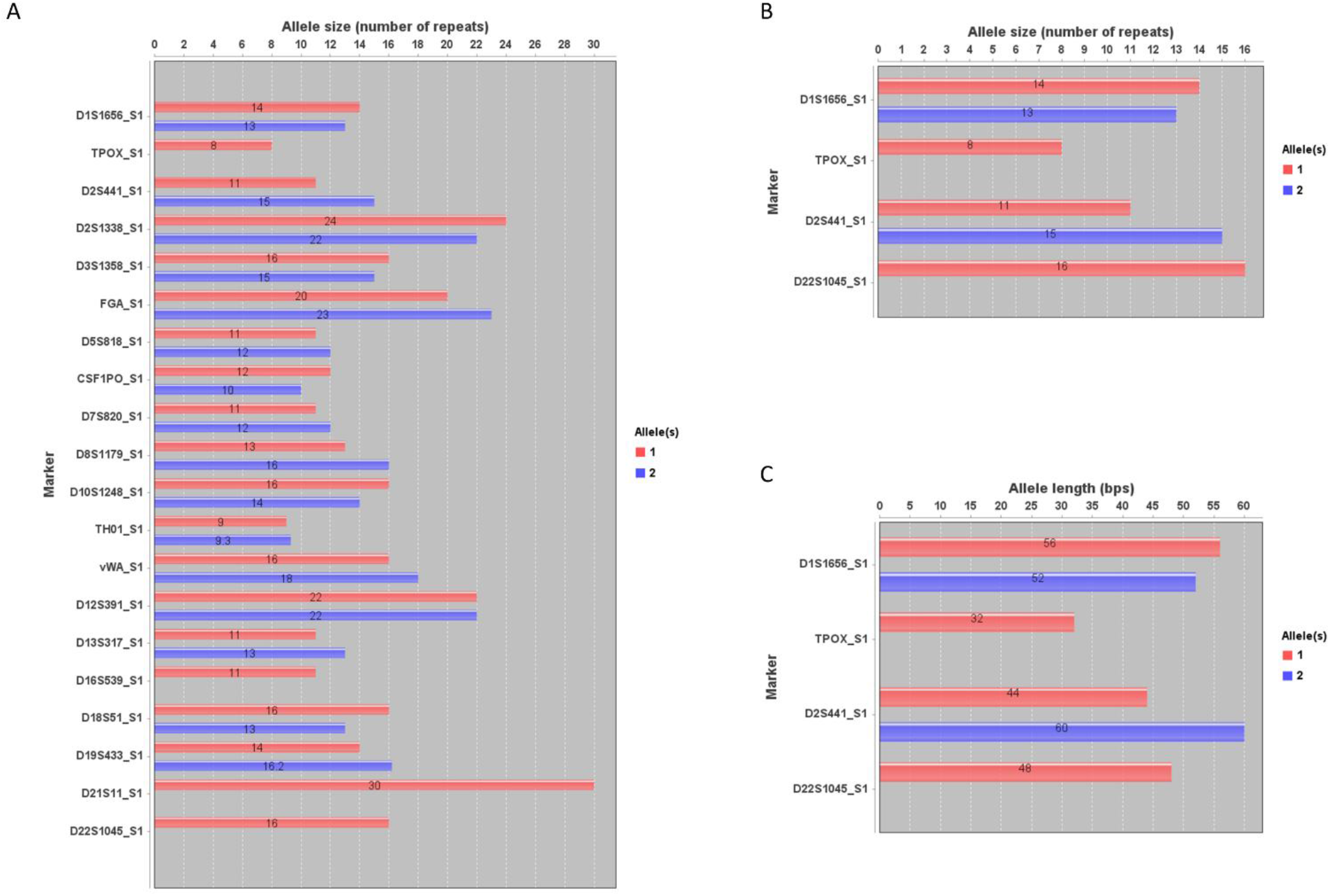
Atlas and distribution plots of tandem repeat alleles. Plots generated by USAT for TR alleles of CODIS core STR markers in the human benchmark sample HG002 from the Genome in a Bottle project. The number on the bar is the detailed size value of the x axis of an allele. A. A bar plot showing the entire atlas of alleles of all 20 CODIS core STR markers. B. A plot showing the comparison of allele sizes of selected markers. C. A plot showing the comparison of allele length in base pairs of selected markers. The x axis shows the number of tandem repeat units or allele length in base pairs. The label next to the y axis shows the name of CODIS STR markers. The callouts in red letters are added annotations for better understanding, not from the software. The name of each bar group is encoded as marker/locus name and sample name joined by an underscore.

### Application of USAT for TR comparison

The above sections demonstrated TR comparisons within a sample or an individual with USAT. To demonstrate the application of USAT for TR comparisons between individuals, TRs from human reference HG002 (son, S1) and HG003 (father, S2) in GIAB were formed as a mixture, and then entered into USAT for analysis. Results of comparing four CODIS STR loci demonstrating a clear TR difference and one copy of TR inheritance between a son and a father are shown in Figure 5 A. To test and visualize the capability of USAT for analyzing mutations, a mutation within the TR allele of CODIS marker allele D1S1656-13 was simulated from sample S1 and marked as S0. The TR sequence comparison successfully showed an expected mutation site in the alignment marked as a dot (Figure 5 B).

**Figure 5.**
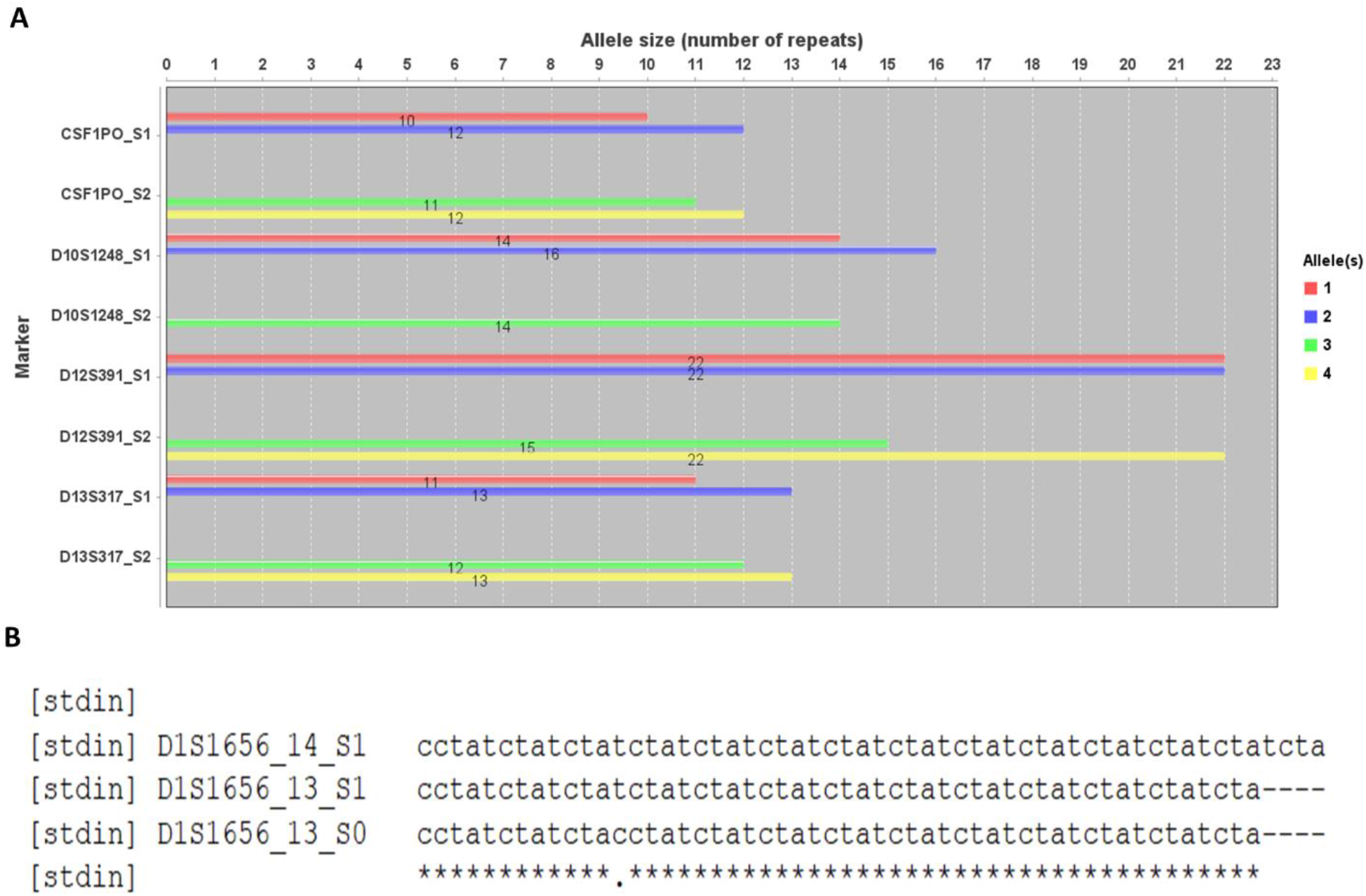
Application example of USAT for TRs between samples. A. An example showing the difference of TR allele sizes between human reference individual HG002 (S1, son) and HG003 (S2, father) from the Genome in a Bottle project. The name of each bar group is encoded as marker/locus name and sample name joined by an underscore. B. An alignment showing the difference between TR haplotypes, where the dot in D1S1656_10_S0 is a simulated mutation in the TR sequence. The name of each sequence is encoded with marker/locus name, allele size, and sample name joined by an underscore.

### Speed

USAT is ultrafast with parallel computing enabled. It took less than one second to analyze 20 CODIS core STR loci of the benchmark human sample HG002 except for the alignment step. The alignment step may take several seconds in Windows 10 system and less than one second in Linux and Mac OS systems.

## Materials and Methods

### TR allele size converting equations

TR size is well defined and used in forensic practice and databases. For applications in forensics, the sequence-based TR alleles can be converted to length-based TR alleles, representing as the number of repeats, for backward compatibility with the forensic DNA databases. The calculation approach is based on the latest recommendation (Alonso *et al*. 2018; Phillips *et al*. 2018), and the alleles were formulated with the equations below. The TR allele sizes usually contain both an integer part and a fractional part, separated by a dot (e.g., 5.1). The fraction part typically is omitted if it is 0.

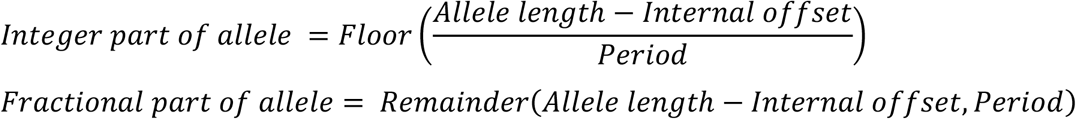

in which the Floor(x) is the function to calculate the greatest integer less than or equal to x; Remainder(x, y) is the remainder of x divided by y; the allele length is the total number of nucleotide bases of an allele; the internal offset is the number of bases that need to be excluded in counting the length-based allele, and the period is the length in base pair of a repeat motif. For example, for a TR with a motif of ATCG (period=4) and an internal offset of 2, the integer part of the allele size of a sequence allele “ATCGATCGggATCGA” (“gg” as internal offset sequences) would be Floor((15-2)/4) = 3, and the factional part is the remainder of (15-2)/4, which is 1, and thus the length-based allele size would be 3.1. To view and compare the haplotype sequences, the length in base pair and the number of repeats are calculated with this formula with the information in the BED file.

### Interactive view and sequence alignment

USAT was programmed with Java JDK 16 (https://www.oracle.com/). JFreeChart Java library (version 1.5.3) (www.jfree.org) was used to plot the figures. To compare the haplotype sequences, the user-selected sequences were dynamically retrieved from the interactive table and entered into MAFFT for a global alignment using Needleman-Wunsch algorithm (Nakamura *et al*. 2018). The alignment is then displayed in an interactive and editable window.

### Testing data

The haplotype sequences of 20 CODIS core STR loci in benchmark human samples HG002 (son) and HG003 (father) were retrieved from the GIAB project. The same sequences were also generated with the ForenSeq kit in a previous study (Gettings et al. 2019). All these sequences were manually verified in a sequence alignment. The haplotype sequences of each STR locus were saved in tabular plain text format according to the required format as an input for USAT (supplementary file).

## Discussion

Genomic sequence variations contain genetic and evolutionay clues of both theoretical and practical value. Increasing amounts of sequence data and studies have enhanced the discovery of sequence-based TR haplotypes (Willems *et al*. 2017; Saini *et al*. 2018; Giesselmann *et al*. 2019; King *et al*. 2021). Further analysis within TR haplotypes could provide additional understanding and interpretation of TR variation and TR’s role in organisms. For example, 25 new sequence variants from 15 CODIS loci were found in an Austrian MPS dataset of 247 reference human samples via sequencing targeted STR loci compared with a length-based CE method (Hölzl-Müller et al. 2021). Such variants were undetectable with traditional CE methods which do not accurately reflect the underlying sequence genotypes. Existing TR associated tools mainly focus on mining TR sequences out of other sequences and report only the length and TR haplotype (e.g., HipSTR (Willems *et al*. 2017)), or the number of repeats and TR haplotype (e.g., STRait Razor (King *et al*. 2021)). Here, our novel USAT software fills the gap for deep TR comparison, comprehensive TR characterization, visualization and comparison of TR sequences.

USAT provides several features for TR applications. The informative TR results generated by USAT are able to facilitate individual identification, TR comparison, marker selection, and further TR marker development (e.g., selecting appropriate TR loci for specific purposes). USAT also provides a direct viewing of the length distribution of TR sequences, which may be used, for example, in TR-associated diagnostic screening of specific diseases. In addition, USAT can provide the detailed descriptions of sequence-based alleles for the STR Sequencing Project (STRSeq) (https://www.ncbi.nlm.nih.gov/bioproject/380127). TRs are widely used for DNA barcoding in many evolution diversity studies, and thus USAT may also be used in biodiversity investigation and discovery of novel species (DeSalle & Goldstein 2019).

More computing threads in general can speed up the analysis in USAT. However, if a dataset is small, two threads in the default setting should be sufficient to obtain results in seconds. In addition, while the alignment of haplotypes may not be in the best format due to multiple possibly acceptable alignments, an additional editing function in the output of alignment is enabled to allow users to adjust the user preferred alignment as needed.

TRs are widely used by a range of researchers with various backgrounds and varying bioinformatic skills. Thus, ease of use for end-users is a very important feature for any bioinformatic tool. USAT was specifically designed for users with limited knowledge and skills in bioinformatics and provides user-friendly graphic interfaces and can be easily adopted by end-users with minimum effort.

## Availability

The USAT software and test data are publicly available at https://github.com/XuewenWangUGA or https://github.com/Ge-Lab

## Supplementary file

USAT_supplementary.docx

