## Supplementary Data for "USAT: a Bioinformatic Toolkit to Facilitate Interpretation and Comparative Visualization of Tandem Repeat Sequences"

UFA: A Software for Tandem Repeat Sequence Interpretation and Visualization

Supplementary data:

### 1. Input data of haplotype for HG002 (S1)

#note: the third sequence for S0 is an simulated sequence with a mutation relative to the $2^{nd}$ sequence.

#CODIS core STR loci for HG002

#Marker_Name Sample_haplotype SampleID

D1S1656 CCTATCTATCTATCTATCTATCTATCTATCTATCTATCTATCTATCTATCTA S1

D1S1656 CCTATCTATCTATCTATCTATCTATCTATCTATCTATCTATCTATCTA S1

D1S1656 CCTATCTATCTACCTATCTATCTATCTATCTATCTATCTATCTA S0

TPOX AATGAATGAATGAATGAATGAATGAATGAATG S1

D2S441 TCTATCTATCTATCTATCTATCTATCTATCTATCTATCTA S1

D2S441 TCTATCTATCTATCTATCTATCTATCTATCTATCTATCTATCTATTTATCTATCTA S1

D2S1338 GGAAGGAAGGACGGAAGGAAGGAAGGAAGGAAGGAAGGAAGGAAGGAAGGAAGGAAGGAAGGAAGGAAGGAAGGCAGGCAGGCAGGCAGGCAGGCAGGCAGGCA S1

D2S1338 GGAAGGAAGGACGGAAGGAAGGAAGGAAGGAAGGAAGGAAGGAAGGAAGGAAGGAAGGAAGGCAGGCAGGCAGGCAGGCAGGCAGGCAGGCAGGCA S1

D3S1358 TCTATCTGTCTATCTATCTATCTATCTATCTATCTATCTATCTATCTATCTATCTATCTA S1

D3S1358 TCTATCTGTCTGTCTATCTATCTATCTATCTATCTATCTATCTATCTATCTATCTA S1

FGA GGAAGGAGAAAGAAAGAAAGAAAGAAAGAAAGAAAGAAAGAAAGAAAGAAAGAAAGAAAGAAAGAGAAAAAAGAAAGAAAGAAA S1

FGA GGAAGGAGAAAGAAAGAAAGAAAGAAAGAAAGAAAGAAAGAAAGAAAGAAAGAAAGAAAGAAAGAAAGAAAGAAAGAGAAAAAAGAAAGAAAGAAA S1

D5S818 ATCTATCTATCTATCTATCTATCTATCTATCTATCTATCT S1

D5S818 ATCTATCTATCTATCTATCTATCTATCTATCTATCTATCTATCT S1

CSF1PO ATCTATCTATCTATCTATCTATCTATCTATCTATCTATCTATCT S1

CSF1PO ATCTATCTATCTATCTATCTATCTATCTATCTATCT S1

D7S820 TATCTATCTATCTATCTATCTATCTATCTATCTATCTATCTATC S1

D7S820 TATCTATCTATCTATCTATCTATCTATCTATCTATCTATCTATCTATC S1

D8S1179 TCTATCTATCTATCTATCTATCTATCTATCTATCTATCTATCTATCTATCTA S1

D8S1179 TCTATCTATCTGTCTATCTATCTATCTATCTATCTATCTATCTATCTATCTATCTATCTATCTA S1

D10S1248 GGAAGGAAGGAAGGAAGGAAGGAAGGAAGGAAGGAAGGAAGGAAGGAAGGAAGGAAGGAA S1

D10S1248 GGAAGGAAGGAAGGAAGGAAGGAAGGAAGGAAGGAAGGAAGGAAGGAAGGAA S1

TH01 AATGAATGAATGAATGAATGAATGAATGAATGAATG S1

TH01 AATGAATGAATGAATGAATGAATGATGAATGAATGAATG S1

vWA TAGATAGATAGATAGATAGATAGATAGATAGATAGATAGATAGACAGACAGACAGACAGATAGA S1

vWA TAGATAGATAGATAGATAGATAGATAGATAGATAGATAGATAGATAGATAGACAGACAGACAGACAGATAGA S1

D12S391 AGATAGATAGATAGATAGATAGATAGATAGATAGATAGATAGATAGATAGATAGACAGACAGACAGACAGACAGACAGACAGAT S1

D12S391 AGATAGATAGATAGATAGATAGATAGATAGATAGATAGATAGATAGATAGATAGACAGACAGACAGACAGACAGACAGACAGAC S1

D13S317 TATCTATCTATCTATCTATCTATCTATCTATCTATCTATCTATC S1

D13S317 TATCTATCTATCTATCTATCTATCTATCTATCTATCTATCTATCTATCTATC S1

D16S539 GATAGATAGATAGATAGATAGATAGATAGATAGATAGATAGATA S1

D18S51 AGAAAGAAAGAAAGAAAGAAAGAAAGAAAGAAAGAAAGAAAGAAAGAAAGAAAGAAAGAAAGAA S1

D18S51 AGAAAGAAAGAAAGAAAGAAAGAAAGAAAGAAAGAAAGAAAGAAAGAAAGAA S1

D19S433 CCTTCCTTCCTTCCTTCCTTCCTTCCTTCCTTCCTTCCTTCCTTCCTTCCTACCTTCTTTCCTT S1

D19S433 CCTTCCTTCCTTCCTTCCTTCCTTCCTTCCTTCCTTCCTTCCTTCCTTCCTTCCTTCCTTCCTACCTTTTCCTT S1

D21S11 TCTATCTATCTATCTATCTGTCTGTCTGTCTGTCTGTCTGTCTATCTATCTATATCTATCTATCTATCATCTATCTATCCATATCTATCTATCTATCTATCTATCTATCTATCTATCTATCTATCTATCTATCTA S1

D22S1045 ATTATTATTATTATTATTATTATTATTATTATTATTATTACTATTATT S1

### 2. Input data of haplotype for HG003

#CODIS core STR loci for HG003(S2)

#Marker_Name Sample_haplotype SampleID

| | | |
|---|---|---|
| D1S1656 | CCTATCTATCTATCTATCTATCTATCTATCTATCTATCTATCTATCTATCTATCTATCTATCTA | S2 |
| D1S1656 | CCTATCTATCTATCTATCTATCTATCTATCTATCTATCTATCTATCTATCTA | S2 |
| TPOX | AATGAATGAATGAATGAATGAATGAATGAATGAATGAATGAATG | S2 |
| TPOX | AATGAATGAATGAATGAATGAATGAATGAATG | S2 |
| D2S441 | TCTATCTATCTATCTATCTATCTATCTATCTATCTATCTATCTA | S2 |
| D2S1338 | GGAAGGAAGGAAGGAAGGAAGGAAGGAAGGAAGGAAGAAAGGAAGGAAGGCAGGCAGGCAGGCAGGCAGGCA | S2 |
| D2S1338 | GGAAGGAAGGACGGAAGGAAGGAAGGAAGGAAGGAAGGAAGGAAGGAAGGAAGGAAGGAAGGAAGGCAGGCAGGCAGGCAGGCA | S2 |
| D3S1358 | TCTATCTGTCTGTCTATCTATCTATCTATCTATCTATCTATCTATCTATCTATCTATCTA | S2 |
| D3S1358 | TCTATCTGTCTATCTATCTATCTATCTATCTATCTATCTATCTATCTATCTATCTATCTATCTA | S2 |
| FGA | GGAAGGAAGGAGAAAGAAAGAAAGAAAGAAAGAAAGAAAGAAAGAAAGAAAGAAAGAAAGAAAGAAAGAAAGAGAAAAAAGAAAGAAAGAAA | S2 |
| FGA | GGAAGGAAGGAGAAAGAAAGAAAGAAAGAAAGAAAGAAAGAAAGAAAGAAAGAAAGAAAGAAAGAAAGAAAGAAAGAAAGAGAAAAAAGAAAGAAAGAAA | S2 |
| D5S818 | ATCTATCTATCTATCTATCTATCTATCTATCTATCTATCTATCT | S2 |
| D5S818 | ATCTATCTATCTATCTATCTATCTATCTATCTATCTATCTATCTATCT | S2 |
| CSF1PO | ATCTATCTATCTATCTATCTATCTATCTATCTATCTATCTATCT | S2 |
| CSF1PO | ATCTATCTATCTATCTATCTATCTATCTATCTATCTATCTATCTATCT | S2 |
| D7S820 | TATCTATCTATCTATCTATCTATCTATCTATCTATCTATC | S2 |
| D7S820 | TATCTATCTATCTATCTATCTATCTATCTATCTATCTATCTATC | S2 |
| D8S1179 | TCTATCTGTCTATCTATCTATCTATCTATCTATCTATCTATCTATCTATCTA | S2 |
| D8S1179 | TCTATCTATCTGTCTATCTATCTATCTATCTATCTATCTATCTATCTATCTATCTATCTATCTA | S2 |
| D10S1248 | GGAAGGAAGGAAGGAAGGAAGGAAGGAAGGAAGGAAGGAAGGAAGGAAGGAA | S2 |
| TH01 | AATGAATGAATGAATGAATGAATGATGAATGAATGAATG | S2 |
| vWA | TAGATAGATAGATAGATAGATAGATAGATAGATAGATAGATAGATAGATAGACAGACAGACAGACAGATAGA | S2 |
| vWA | TAGATAGATAGATAGATAGATAGATAGATAGATAGATAGATAGATAGACAGACAGACAGACAGATAGA | S2 |
| D12S391 | AGATAGATAGATAGATAGATAGATAGATAGATAGATAGATAGATAGATAGATAGATAGACAGACAGACAGACAGACAGACAGACAGACAGAC | S2 |
| D12S391 | AGATAGATAGATAGATAGATAGATAGATAGATAGACAGACAGACAGACAGACAGACAGAT | S2 |

D13S317 TATCTATCTATCTATCTATCTATCTATCTATCTATCTATCTATCTATC S2

D13S317 TATCTATCTATCTATCTATCTATCTATCTATCTATCTATCTATCTATCTATC S2

D16S539 GATAGATAGATAGATAGATAGATAGATAGATAGATAGATAGATA S2

D16S539 GATAGATAGATAGATAGATAGATAGATAGATAGATA S2

D18S51 AGAAAGAAAGAAAGAAAGAAAGAAAGAAAGAAAGAAAGAAAGAAAGAAAGAAAGAAAGAA S2

D18S51 AGAAAGAAAGAAAGAAAGAAAGAAAGAAAGAAAGAAAGAAAGAAAGAAAGAAAGAA S2

D19S433 CCTTCCTTCCTTCCTTCCTTCCTTCCTTCCTTCCTTCCTTCCTTCCTACCTTCTTTCCTT S2

D21S11 TCTATCTATCTATCTATCTGTCTGTCTGTCTGTCTGTCTGTCTATCTATCTATATCTATCTATCTATCATCTATCTATCCATATCTATCTATCTATCTATCTATCTATCTATCTATCTATCTATCTATCTA S2

D21S11 TCTATCTATCTATCTATCTGTCTGTCTGTCTGTCTGTCTGTCTATCTATCTATATCTATCTATCTATCATCTATCTATCCATATCTATCTATCTATCTATCTATCTATCTATCTATCTATCTA S2

D22S1045 ATTATTATTATTATTATTATTATTATTATTATTATTATTACTATTATT S2

D22S1045 ATTATTATTATTATTATTATTATTATTATTATTATTACTATTATT S2
